# The 5-Hydroxytryptaminergic Neurons in the Dorsal Raphe Nucleus and Different 5-HT Receptors are Implicated in Mediating the Awakening from Sevoflurane Anesthesia

**DOI:** 10.1101/2022.05.22.492958

**Authors:** HaiXiang Ma, LeYuan Gu, YuLing Wang, Qing Xu, Qian Yu, XiTing Lian, WeiHui Shao, Lu Liu, JiaXuan Gu, Yue Shen, HongHai Zhang

## Abstract

In order to explore the mechanism of general anesthesia emergence, based on the common clinical phenomenon-delayed emergence, we explore the role of 5-hydroxytryptamine (5-HT) neurons in the dorsal raphe nucleus in promoting awakening from sevoflurane anesthesia in mice model. In this study, C57BL/6J male mice were selected to specifically activate or inhibit 5-HT neurons in the dorsal raphe nucleus (DRN) and different 5-HT receptors by intraperitoneal, lateral ventricle, intranuclear or DRN injection of agonists/antagonists and optogenetics during the sevoflurane anesthesia to record and observe the anesthesia induction and emergence time of mice. Through intraventricular infusion and intranuclear microinjection of 5-HT and the agonists or antagonists of different 5-HT receptors, our data showed that 5-HT and 5-HT1A and 2A/C receptors, especially 5-HT1A receptor, are involved in the regulation of delayed awakening mediated by DRN 5-HT neurons. This can provide a reliable theoretical basis as well as potential targets for clinical intervention to prevent delayed emergence and some postoperative risks.

**Graphical Abstract:** 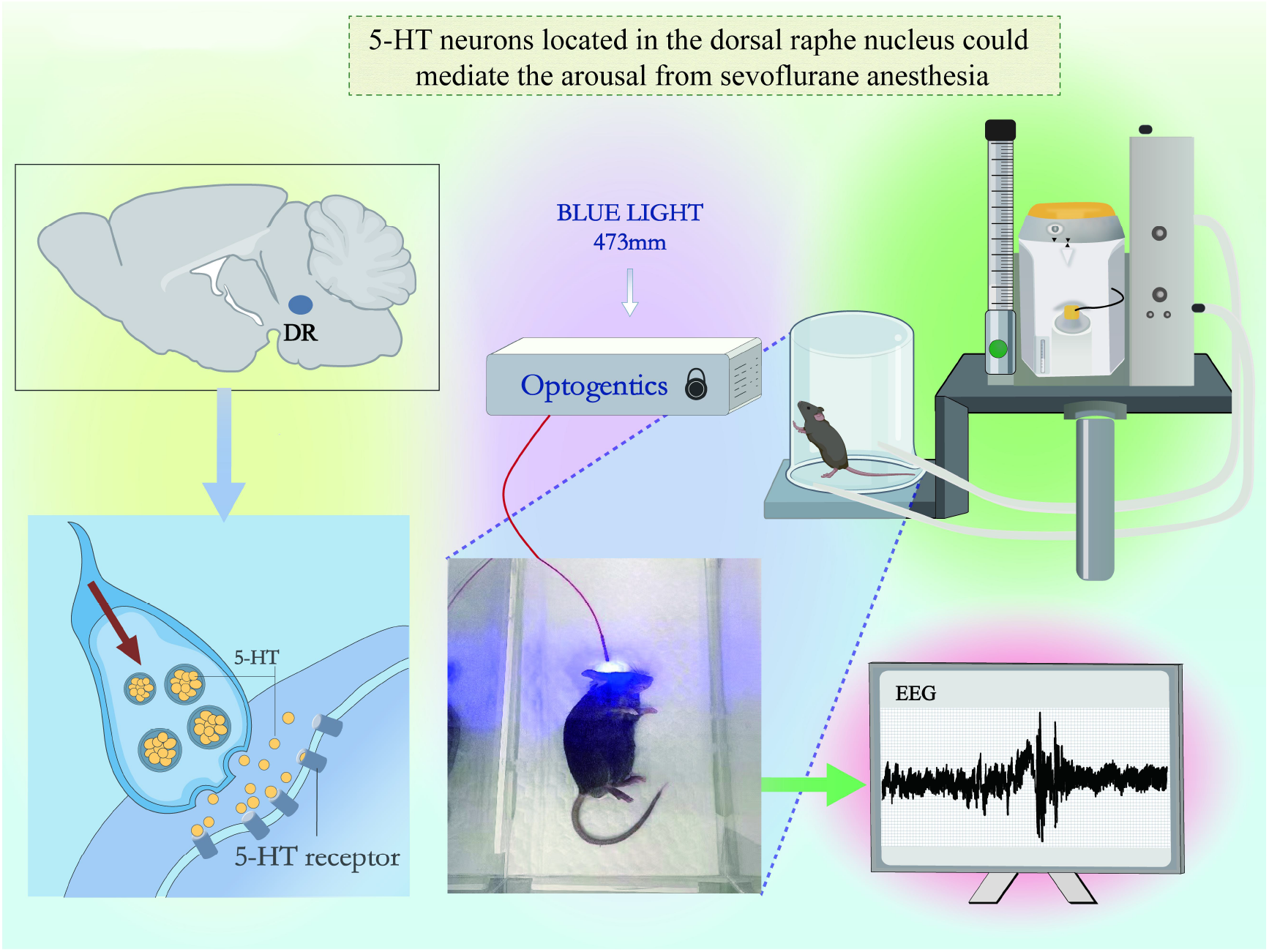

## Introduction

General anesthesia is widely used in surgery or a variety of diagnostic medicine due to its effect of reversible loss of consciousness and recovery of consciousness^[1–4]^. Inhaled sevoflurane is one of the most prevalent methods of general anesthesia, and can be used for induction as well as maintenance of general anesthesia. General anesthesia can cause some serious complications[5], such as delayed emergence, which might lead to an increased incidence of cardiovascular accidents[6], reduced cognition level[7], and a variety of other central and peripheral complications[8–11]. However, the mechanisms of delayed emergence remain elusive.

Several studies have reported that 5-HT levels could fluctuate during anesthesia, further indicating the regulatory role of the 5-HT system in general anesthesia[12,13]. However, how 5-HT neurons and 5-HT receptors regulate general anesthesia has been poorly studied. 5-HT, as an important neurotransmitter, is involved in many functions of the central system, including sleep-wake behavior, cognition, emotion, sexual function, thermoregulation and food intake[14,15]. 5-HT in the central nervous system is mainly located in the dorsal raphe nucleus (DRN). The 5-HT neurons of DRN mainly project to the midbrain and forebrain, predominately the cerebral cortex, limbic system, basal forebrain, and hypothalamus[16–18]. Numerous studies have demonstrated the importance of 5-HT in the sleep-wake cycle[19,20], suggesting that the 5-HT neuron system may be crucial in mediating arousal. Given that the delayed emergence caused by general anesthesia is closely related to the arousal function of the nervous system, while 5-hydroxytryptamine (5-HT) is involved in regulating arousal, we focused our attention on the 5-HT system. And explaining the important nuclei associated with sleep-wake regulation, especially the 5-HT neurons of DRN, is particularly necessary for studying the regulation of general anesthesia awakening.

In the present research, we performed pharmacological experiments and photogenetic experiments to regulate the concentration of 5-HT in different levels and the activity of 5-HT neurons in the nucleus. We aim to explore whether exogenous 5-HT supplementation or manipulation of the 5-HT neurons of DRN could affect recovery time under sevoflurane anesthesia. Finally, pharmacological methods were used to explore the role of specific 5-HT receptor subtypes in anesthesia awakening.

Our results showed that 5-HT neurons in DRN played a key role in sevoflurane anesthesia. Activating 5-HT neurons in DRN can reduce the emergence time, and various kinds of 5-HT receptors have different effects on the recovery from sevoflurane anesthesia. This study can serve as a scientific foundation for investigating the central mechanism of general anesthesia and clinical regulation of post-anesthesia recovery phase. Most importantly, it may provide a potential target for intervention in preventing delayed awakening from general anesthesia.

## Materials and Methods

### 2.1 Animals

The experimental animals in this study were wild-type mice (C57BL/6J), purchased from the Animal Experiment Center of Zhejiang University and approved by the Animal Advisory Committee of Zhejiang University. All mice were housed and bred in the SPF-Class Housing of Laboratory of Animal Center of Zhejiang University School of Medicine and given rodent food and water ad libitum. The indoor temperature was 25°C, humidity was 65%, and light cycle was maintained for 12 hours. All experimental procedures were in line with the National Institutes of Health Guidelines for the Care and Use of Laboratory Animals and approved by the Animal Advisory Committee of Zhejiang University. In order to avoid the interference of gender and female estrous cycle, only male mice were used, and all the experiments were completed between 9:00 and 15:00.

### 2.2 Pharmacological experiments

#### 2.2.1 Effect of IP administration of 5-HTP on induction time and emergence time of sevoflurane anesthesia

For intraperitoneally (IP) administration, 5-HTP was dissolved in saline. Intraperitoneal injections of 5-HTP (50 mg/kg, 100 mg/kg), a precursor to 5-HT synthesis, were administered to C57BL/6J mice, and 24 h later, 5-HTP was injected intraperitoneally again. One hour later, the mice were placed in an anesthesia box for induction and maintenance, and induction time was recorded, ceasing the anesthesia 30 minutes after the start of anesthesia and record the emergence time.

#### 2.2.2 Effect of ICV administration of different agonists and antagonists of 5-HT receptors on induction time and emergence time of sevoflurane anesthesia

The same batch of 8-week-old wild-type C57BL/6J mice with unilateral ventricular catheter implantation for 1 week was used. The mice were administered agonists/antagonists of different 5-HT receptors through the lateral ventricle and placed in an anesthesia box for anesthesia induction and maintenance, and the anesthesia induction time was recorded. 5-HT1A receptor agonist 8-OH-DAPT (3mg/ml, 6mg/ml) or 5-HT1A receptor antagonist WAY 100635 (5mg/ml, 10mg/ml) or 5-HT2A/C receptor agonist DOI (1mg/ml, 2mg/ml) or 5-HT2A receptor antagonist KET (Ketanserin) (1mg/ml, 2mg/ml) 2000nL were administered through a lateral ventricular catheter at 15 minutes of anesthesia. The anesthesia was stopped at 30 minutes and the emergence time was recorded.

#### 2.2.3 Effect of DRN administration of different agonists and antagonists of 5-HT receptors on induction time and emergence time of sevoflurane anesthesia

The same batch of 8-week-old wild-type C57BL/6J mice with DRN guide cannula implantation for 1 week was used and was microinjected agonists/antagonists of different 5-HT receptors into the nucleus through cannulas. C57BL/6J mice were placed in an anesthesia box for anesthesia induction and maintenance, and the anesthesia induction time was recorded as induction time. After the 15 minute of anesthesia, 8-OH-DAPT (1.5mg/ml, 3mg/ml, 6mg/ml) or WAY 100635 (5mg/ml, 10mg/ml, 15mg/ml) or DOI (1mg/ml, 2mg/ml) or KET (1mg/ml, 2mg/ml) 200nL was administered through the cannulas. The anesthesia was stopped 30 minutes after anesthesia and the time of awakening was recorded as emergence time.

### 2.3 Stereotactic localization and virus injection

#### 2.3.1 Stereotactic surgery

C57BL/6J mice were anesthetized with 3.5% chloral hydrate and head-fixed in a stereotaxic apparatus (68030, RWD Life Science Inc., Shenzhen, China), as previously described. Throughout the entire surgical process, the body temperature of anesthetized mice was kept constant at 37°C using a heating pad. According to the 4th edition of mouse brain atlas, the target brain coordinates were located on the skull surface (DRN: AP−4.60mm, ML−0.05 mm, DV−3.10 mm, 10° right; parococele: AP−0.45 mm, ML−1.00 mm, DV−2.50 mm). After the coordinates were confirmed, a miniature hand-held cranial drill penetrated the skull without damaging the dura mater and cerebral parenchyma. Bone fragments and blood scabs were removed with the tip of a 1ml syringe. For optogenetic viral delivery of pAAV-TPH2 PRO-ChETA-EYFP-WPRES-PAS (1013vg/ml), which was purchased from Sheng BO, Co., Ltd. (Shanghai, China) and the sequences of vectors were designed by Kazuki Nagayasu as the gift to the study (Department of Molecular Pharmacology Graduate School of Pharmaceutical Sciences, Kyoto University), microinjection (100 nl, 40 nl/min) was performed via a gauge needle by an Ultra Micro Pump (160494 F10E, WPI) over a period of 10 min; the syringe was not removed until 15 min after the end of infusion to allow diffusion of the viruses. Wet cotton balls with normal saline are placed on the surface of the skull to keep the skull and scalp moist, helping to ease skin tension and facilitate suturing. The incision was smeared with erythromycin ointment to avoid infection. Then the mice were resuscitated on the thermostatic electric blanket, and put back into the feeding cage after confirming reactivation. Wait until the virus has been infected for 3-4 weeks before proceeding to the next step. After the experiment, the position of virus expression and cannula was confirmed, that is, whether the fluorescence expression of the virus and cannula was buried in the target brain region.

#### 2.3.2 Lateral ventricle catheterization (ICV) and electroencephalogram (EEG) electrode implantation

After leveling the skull and drilling the target region, a shallow concave is drilled in the left anterior bregma and the left and right anterior of the posterior fontanelle respectively (without drilling through the skull). Fix the screw in a shallow concave, with exposed 1/3 of the skull surface and firmly connected as the standard. To embed electrodes of the EEG, a wire is fixed to the screws in the left anterior bregma and the right anterior of the posterior fontanelle, and then tighten the screws, so that the top of the screw contacts the brain tissue, and then secures the cannula and screws. After that, two wires were welded to the electrode. Then, the cannula, screws, electrodes, and skull were tightly cemented and inserted into the inner core of the cannula, and the mice were weighed again and recorded. If electrode embedding is not required, the cannula is directly embedded into the target brain area. Erythromycin ointment was applied to the surgical area and the mice were fed alone to recover for one week.

#### 2.3.3 Photometric analysis of induction time and emergence time of sevoflurane anesthesia in DRN

pAAV-TPH2 PRO-ChETA-EYFP-WPRES-PAS was delivered into the DRN of C57BL/6J mice, and three weeks after virus injection, optical fiber (diameter 200μm, numerical aperture 0.22) was implanted in the DRN. After the fiber was embedded for one week, the optogenetics experiments were performed, and the laser was connected to the core of the mice’s heads by the optical fiber bouncing lining. The induction time and emergence time of sevoflurane anesthesia were analyzed statistically. The 5-HT neurons (expressing ChETA) in DRN were activated by blue light. The parameters for photostimulation of the DRN were: blue-light, 465 nm, 20 Hz, 20-ms pulse width, 15 mW, and 20 min. Adjust the output parameters of the laser, detect the laser intensity of the end of the core with the optical power meter, and control the laser range within 15mW.

### 2.4 Behavioral tests

Behavioral tests were adopted to examine the induction and emergence times. Mice were freely placed in an inhalation anesthesia box for 30 min to adapt to the experimental environment. The gas flow into the anesthesia box is 2L/min, with 80% oxygen and 20% nitrogen. Before the experiment, the bottom of the anesthesia box was preheated with an electric heating pad for 15min to keep the temperature in the anesthesia box appropriate during the experiment. Soda lime was spread on the bottom of the box to absorb the CO2. Then, anesthesia was induced by 8MAC sevoflurane with 100% O2 at a flow rate of 2 L/min. The box was rotated 90 degrees every 15s until the mice exhibited loss of righting reflex (LORR) and could not turn themselves prone onto all four limbs. The time from the initiation of sevoflurane inhalation to LORR was considered as induction time. At the 30 th minute of anesthesia, the sevoflurane was interrupted by shutting down, and the residual sevoflurane in the anesthesia box and tube was removed by rapid oxygen administration at a rate of 2 L/min. The anesthesia box was rotated as described above to evaluate the time when mice could independently turn from the supine position with at least three paws reaching the bottom of the box. The time from the cessation of anesthesia to the recovery of righting reflex (RORR) was defined as emergence time.

### 2.5 Immunohistochemistry and histology

The position of the optical fiber cannula tip for the DRN in each mouse was verified by histology. After approximately 90 min in the experiment, C57BL/6J mice were sacrificed and perfused with phosphate-buffered saline (PBS) containing 4% paraformaldehyde (PFA). After fixation in 4% PFA for about 16h and saturation in 30% sucrose (24h), each mouse brain was sectioned into 30-μm-thick coronal slices with a freezing microtome (CM30503, Leica Biosystems, Buffalo Grove, IL, USA). The sections were first washed in PBS three times for 5 min each and incubated in blocking solution for 2.5h at room temperature. Then for c-fos or TPH2 staining in the DRN, sections were incubated at 4°C overnight in a solution of mouse anti-c-fos primary antibody (1:1000 dilution, Abcam, UK) or mouse anti-TPH2 primary antibody (1:100 dilution, T0678, Sigma-Aldrich, UK). The secondary antibodies used were anti-mouse Alexa 488 (1:1000; A32766, Thermo Fisher Scientific, Waltham, MA, USA), anti-rabbit Alexa 546 (1:1000; A10036, Thermo Fisher Scientific), or anti-rabbit Alexa 488 (1:1000; A10523, Thermo Fisher Scientific), and slices were incubated in secondary antibody solutions for 1h at room temperature. After washing with PBS three times for 15 min each, the sections were mounted onto glass slides and incubated in DAPI (1:4000; Cat#C1002; Beyotime Biotechnology; Shanghai, China) for 7 min at room temperature. Finally, the glass slides were sealed using 60% glycerol. All images were taken with a Nikon A1 laser-scanning confocal microscope (Nikon, Tokyo, Japan). The numbers of immunopositive cells were counted and analyzed using ImageJ (NIH, Bethesda, MD, USA). Notably, mice in which the implantation placement was outside of the targeted brain structure were not used in our experiments. Positively stained cells co-expressing c-fos, and TPH2 were counted as previously described.

### 2.6 Statistical analysis

All data are presented as the mean ± standard error of the mean (SEM). Statistical analyses were performed using GraphPad Prism TM 8.0 and SPSS version 22.0 (SPSS Software Inc., Chicago, IL, USA). Two groups of comparisons: If the data is normally distributed, use the Student’s T test, including independent-samples T test and paired-samples T test. If the data is not normally distributed, use Mann-Whitney U or Wilcoxon signed-rank test. Levene test was used to test the homoscedasticity of each other. After the data is in line with normal distribution and homoscedasticity, one-way ANOVA was used for the comparison of three groups and above, and two-way ANOVA was used for the comparison of double-factor, followed by Bonferroni’s test. Statistical significance was inferred if P<0.05.

## Results

### 3.1 IP injection of 5-HTP reduced induction and emergence time of sevoflurane anesthesia

To explore whether and how 5-HT affects sevoflurane anesthesia induction and emergence time, we selected wild-type C57 mice at 8 weeks after birth for the experiment. C57 mice were intraperitoneally injected 5-HTP at different doses (50, 100mg/kg), the precursor of 5-HT, 24 hours after the injection, again intraperitoneally injected 5-HTP. After 1h, the mice were placed in an anesthesia box and 8MAC sevoflurane for induction. When the mice exhibited LORR and could not turn themselves prone onto all four limbs, recorded as induction time and changed to 2MAC sevoflurane for anesthesia maintenance. The inhalation of sevoflurane was stopped at the 30th min, and rapid oxygen delivery at a rate of 2 L/min was used to quickly clean the residual sevoflurane. The time from the cessation of anesthesia to the RORR was defined as emergence time. Compared with the control group, IP injection of 5-HTP (100mg/kg) prolonged induction time (P<0.05) and reduced emergence time(P<0.01, Figure 1).

**Figure 1.**
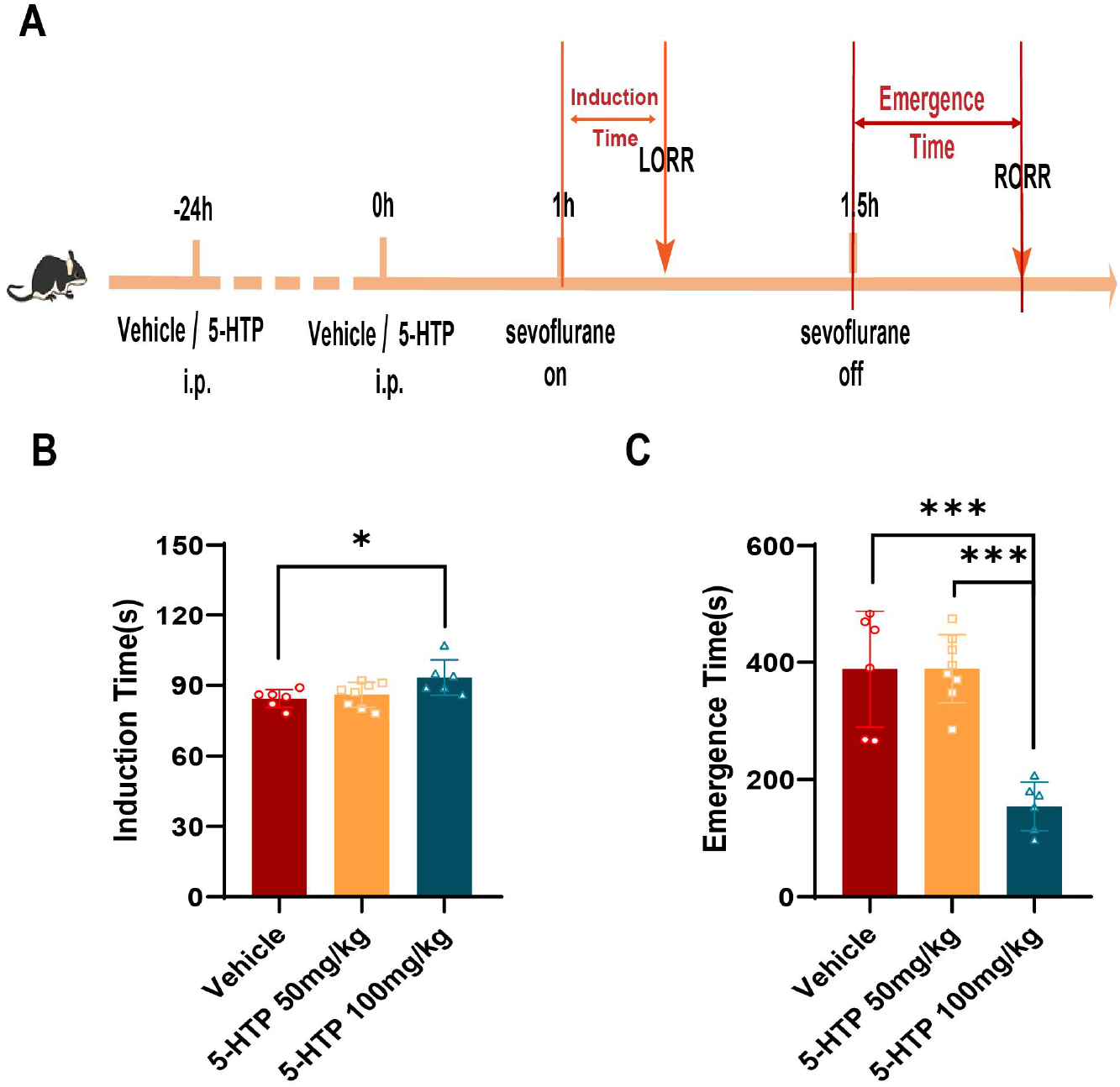
Effect of intraperitoneal injection of 5-HTP on sevoflurane anesthesia induction time and emergence time. A. Experimental process of IP injection of 5-HTP. B. Induction time of sevoflurane anesthesia in vehicle (NS) and different doses of 5-HTP group. Compared with the control group (n=6) and 50mg/kg 5-HTP group (n=8), IP administration of 100mg/kg 5-HTP group had a longer induction time (n=6, P<0.05). C. Emergence time of sevoflurane anesthesia in vehicle and different doses of 5-HTP group. Compared with the control group (n=6) and 50mg/kg 5-HTP group (n=8), IP administration of 100mg/kg 5-HTP group held a shorter emergence time (n=6, P<0.05), while there was no significant difference between the 50mg/kg 5-HTP group and the vehicle group.

### 3.2 The activity of 5-HT neurons in the DRN was significantly reduced by the sevoflurane anesthesia

To investigate the activity of 5-HT neurons in the DRN under sevoflurane anesthesia, we observed the changes of the co-expression between the c-fos and TPH2 in DRN by immunohistochemistry. Compared with the control group without the sevoflurane anesthesia group, the co-expression of c-fos and TPH2 was significantly reduced in the group with the sevoflurane anesthesia, which suggested that the activity of 5-HT neurons in the DRN can be suppressed by sevoflurane anesthesia (P<0.05, Figure 2). It may be a major cause of delayed emergence.

**Figure 2.**
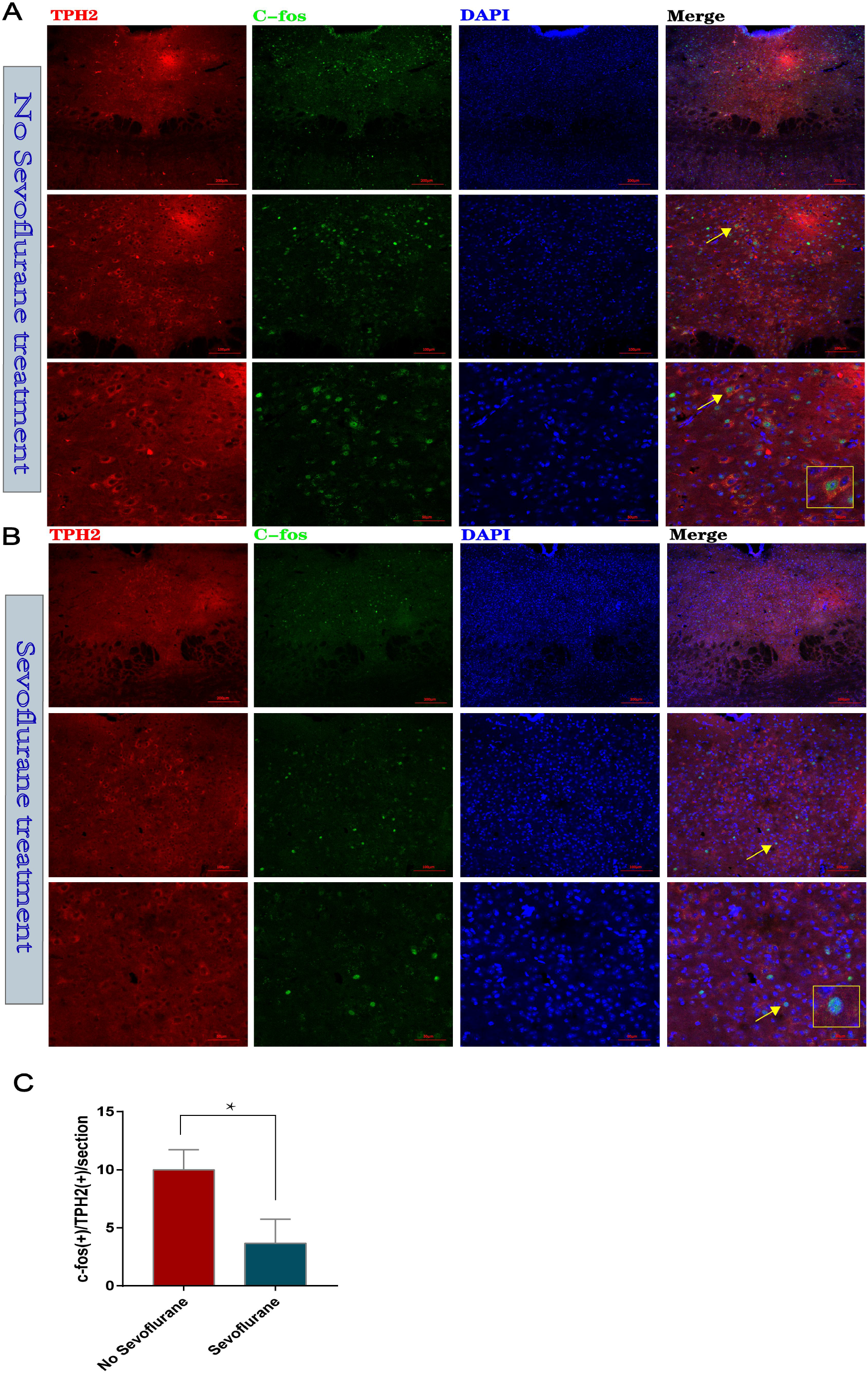
Effect of the sevoflurane anesthesia on the co-expression of c-fos and TPH2 in DRN. A-C. Compared with the control group without treatment with the sevoflurane anesthesia (n=3), the co-expression of c-fos and TPH2 in DRN was significantly reduced in the group with the sevoflurane anesthesia (n=3/group, P<0.05).

### 3.3 Activation of 5-HT neurons in the DRN reduced the emergence time of sevoflurane anesthesia

The experiment of the first part showed that the nonspecific supplement of exogenous 5-HTP to increase 5-HT could reduce the emergence time of sevoflurane anesthesia. To further explore the effects of 5-HT neurons in DRN on induction time and emergence time of sevoflurane anesthesia, we selectively enhanced 5-HT neurotransmission on the DRN by applying photostimulation (blue light, 15mW, 20Hz). ChETA with TPH2 was injected into the DRN of 8-week-old mice to induce specific expression of the virus in 5-HT neurons. Fiber implantation was performed at the DRN 3 weeks later, and the experiment was performed 1 week after surgery. The mice were placed in an anesthesia box with 8MAC sevoflurane for induction. When the mice exhibited LORR and could not turn themselves prone onto all four limbs, this was recorded as induction time and changed to 2MAC sevoflurane for anesthesia maintenance. At the 10th minute of anesthesia, the experimental group performed photostimulation (PS) with blue light at 465 nm (15 mW,20 Hz) to activate 5-HT neurons in the DRN, while the control group did not perform photostimulation (No PS). The inhalation of sevoflurane was stopped at the 30th min, and rapid oxygen delivery at a rate of 2 L/min was used to quickly clean the residual sevoflurane. The time from the cessation of anesthesia to the RORR was defined as emergence time (Figure 3 A). Compared with the control group, there was no significant difference in the induction time of mice in each experimental group (P>0.05, Figure 3 B). Both 15min and 20min blue light irradiation at the DRN significantly reduced the emergence time (P<0.01, Figure 3 B), while 5min blue light irradiation showed no significant difference in the emergence time (P>0.05, Figure 4 C). These data indicated that specific activation of 5-HT neurons in DRN can significantly reduce the emergence time, but has no significant effect on induction time.

**Figure 3.**
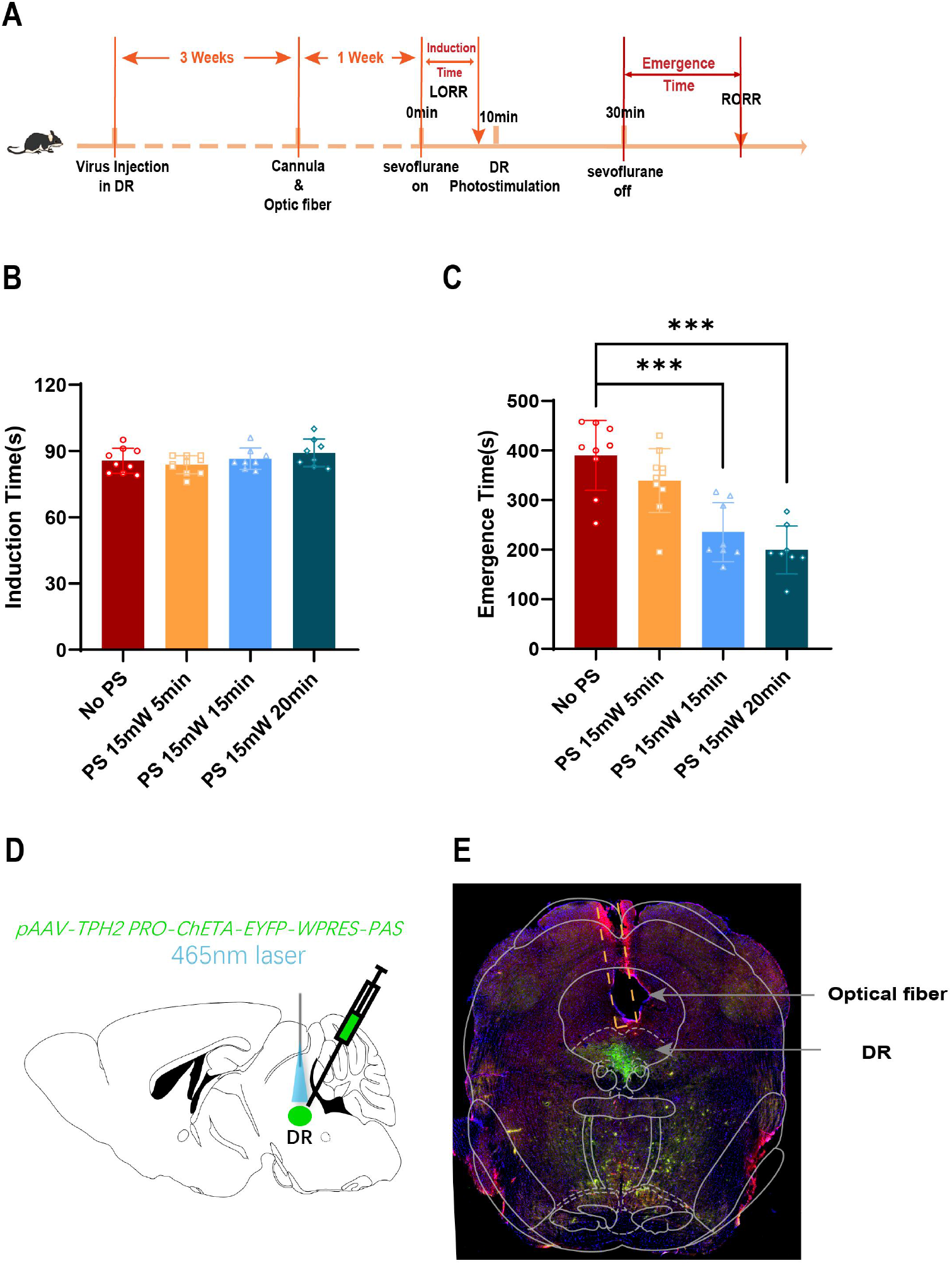
Effect of 5-HT neurons activated by optogenetics on sevoflurane induction time and emergence time in DRN. Experimental process of activation of 5-HT neurons in DRN by optogenetics. B. Induction time in control group without light and group with different light parameters. Compared with the control group (No PS, n=9), there was no significant difference in the induction time of mice in each experimental group (PS 15mW 5min: n=10, PS 15mW 15min: n=8, PS 15mW 20min: n=8, P>0.05). C. Emergence time in control group without light and group with different light parameters. Compared with the control group (No PS, n=9), emergence time of PS 15mW 15min group and 20min group was significantly decreased (PS 15mW 15min: n=8, PS 15mW 20min: n=8, P<0.01), while the awakening time of PS 5mW 5min (n=10) group was no different from the control group. Data are mean±SEM. D. Schematic diagram of ChETA virus injection in DRN and optical fiber location and colocalization with TPH2. E. Immunohistochemistry of C57 mice showed the location of fiber tip and revealed co-expression neurons of ChETA and TPH2. Thermal damage induced by light stimulation was not observed in the region around the fiber tip.

**Figure 4.**
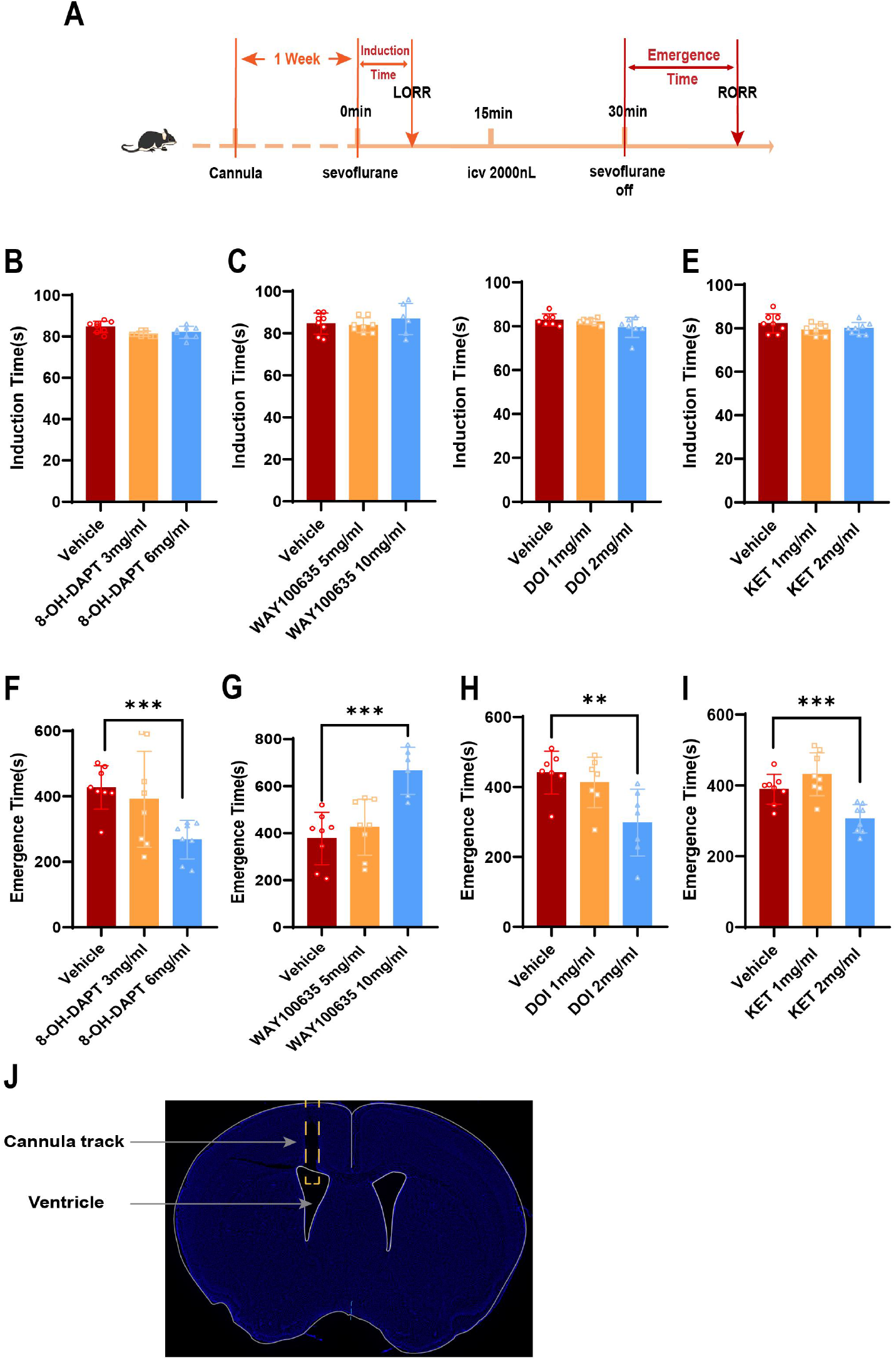
Effects of different 5-HT agonists/antagonists injected into the lateral ventricle (ICV) on induction time and emergence time of sevoflurane anesthesia. Specific experimental protocol for lateral ventricle administration. (B-E) Induction time of sevoflurane anesthesia in different concentrations of 8-OH-DAPT, WAY 100635, DOI and KET groups; There was no statistical difference in induction time among all groups (P>0.05, n=8). (F-I) Emergence time of sevoflurane anesthesia in groups with different concentrations of 8-OH-DAPT, WAY 100635, DOI and KET (n=8). (F) Compared with the Vehicle group, the emergence time of 6mg/mL 8-OH-DAPT group was significantly reduced (P<0.01). (G) Compared with Vehicle, the emergence time of 10mg/mL WAY 100635 group was significantly reduced (P<0.01). (H) compared with the Vehicle group, the emergence time of the 2mg/ml DOI group was significantly reduced (0.01<P<0.05). (I) compared with the Vehicle group, the emergence time of the 2mg/ml KET group was significantly reduced (P<0.01). (J) Staining results of the position of the lateral ventricular cannula; Data are mean±SEM.

After the experiment, immunohistochemistry was performed to determine the position of virus injection and optical fiber embedding. TPH2 and ChETA were co-located in the DRN region of mice brains to determine the location of virus injection (Figure 3 D) and the colocalization of virus and 5-HT neurons (Figure 3 E).

### 3.4 Effects of different 5-HT agonists/antagonists injected into the lateral ventricle (ICV) on induction time and emergence time of sevoflurane anesthesia

To explore the effects of different 5-HT receptors on anesthesia induction and emergence time, we selected wild-type 8-week-old C57 mice for lateral ventricle catheterization and administered agonists/antagonists of different 5-HT receptors by ICV. After cannula implantation one week, C57 mice were placed in an anesthesia box with 8MAC sevoflurane for induction. When the mice exhibited LORR and could not turn themselves prone onto all four limbs, recorded as induction time and changed to 2MAC sevoflurane for anesthesia maintenance. At 15 th minute of anesthesia, 5-HT1A receptor agonist 8-OH-DAPT, 5-HT1A receptor antagonist WAY100635, 5-HT2A/C receptor agonist DOI or 5-HT2A receptor antagonist KET 2000nL was administered through a lateral ventricle tube. Compared with the control group (vehicle), the induction time of sevoflurane anesthesia was not significantly affected by the administration of 8-OH-DAPT, WAY100635, DOI or KET in the DRN (Figure 4 B-E). However, in terms of the time to emergence, compared with the control group, administration of 8-OH-DAPT (6mg/ml) as well as DOI (2mg/ml) or KET (2mg/ml) in DRN nucleus reduced the emergence time (P<0.01 or 0.01<P<0.05, Figure 4 F, H, I), while administration of WAY100635 (10mg/ml) in DRN nucleus increased the emergence time (P<0.01, Figure 4 G). And administration of 8-OH-DAPT and KET significantly shorten the awakening time (P<0.01, Figure 5 F, I). These results showed that activation of 5-HT1A receptor could significantly shorten the time of recovery from sevoflurane anesthesia, while antagonism of 5-HT1A receptor could significantly prolong the time to recovery from sevoflurane anesthesia. However, for 5-HT2A and C receptors, antagonizing 5-HT2A receptor can significantly shorten the emergence time, but simultaneous activation of 5-HT2A and C receptors can also shorten the emergence time, but not as obviously as antagonizing 5-HT2A alone. In conclusion, different 5-HT receptors may play different roles in the delay of awakening. Combined with the previous results of intraperitoneal injection of 5-HTP and activation of 5-HT neurons in DR by optogenetics, we speculated that the 5-HT1A receptor may play a major role in awakening under general anesthesia.

**Figure 5.**
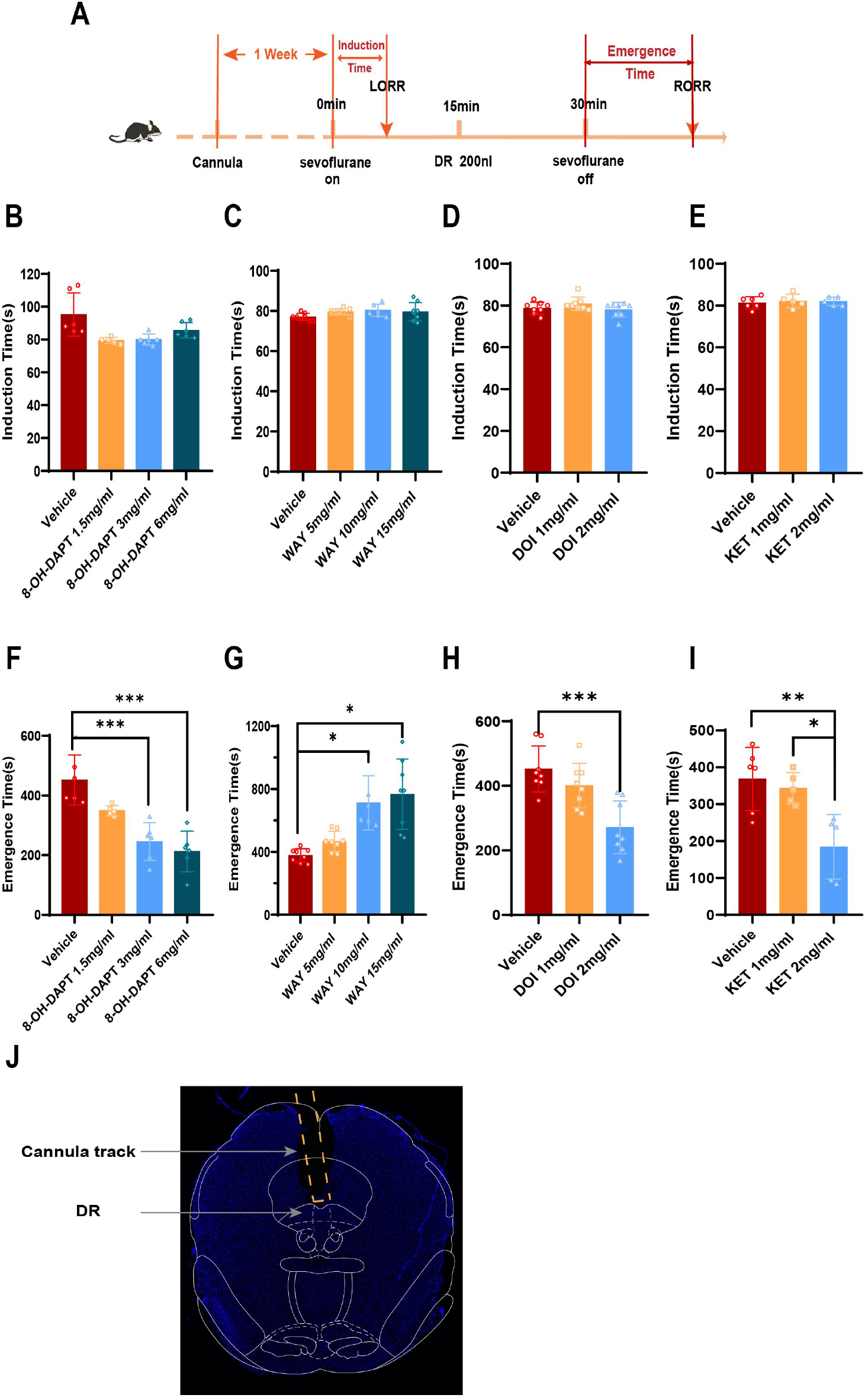
Effect of 5-HT agonist/antagonist microinjection in DR nucleus on induction and recovery time of sevoflurane anesthesia. Specific experimental protocol for intranuclear microinjection of DR. (B-E) Induction time of sevoflurane anesthesia in different concentrations of 8-OH-DAPT, WAY 100635, DOI and KET groups; There was no statistical difference in induction time among all groups (P>0.05, n=8). (F-I) Emergence time of sevoflurane anesthesia in groups with different concentrations of 8-OH-DAPT, WAY 100635, DOI and KET (n=8). (F) Compared with the Vehicle group, the emergence time of 3mg/mL and 6mg/mL 8-OH-DAPT groups was significantly reduced (P<0.01). (G) Compared with Vehicle, the emergence time of 10mg/mL and 15mg/mL WAY 100635 group was significantly decreased (P<0.05). (H) Compared with the vehicle group, the emergence time of the 2mg/ml DOI group was significantly reduced (P<0.01). (I) Compared with the vehicle group, the emergence time of the 2mg/ml KET group was significantly reduced (P<0.05). (J) Staining results of the position of the DR cannula, Data are mean±SD.

### 3.5 Effect of 5-HT agonist/antagonist microinjection in DRN on induction and emergence time of sevoflurane anesthesia

The fourth part of the experiment showed that different 5-HT receptors in the central nervous system had different effects on the emergence time of sevoflurane anesthesia. In order to further clarify the effects of different 5-HT receptors in DRN on sevoflurane anesthesia induction and emergence time in mice, the same batch of 8-week-old wild-type C57BL/6J mice with DRN guide cannula implantation for 1 week was used and was microinjected agonists/antagonists of different 5-HT receptors into the nucleus through cannulas. C57BL/6J mice were placed in an anesthesia box for anesthesia induction and maintenance one week after DR cannula insertion, and the anesthesia induction time was recorded. At the 15 th minute of anesthesia, 8-OH-DAPT or WAY 100635 or DOI or KET 200nL was administered through the cannulas. The anesthesia was stopped 30 minutes after anesthesia and the time of awakening was recorded (Figure 5 A). The experimental results showed that compared with the vehicle group, intra-DRN microinjection of 8-OH-DAPT or WAY 100635 or DOI or KET had no significant effect on sevoflurane induction time (P>0.05, Figure 5 B-E). However, for emergence time, microinjection of 8-OH-DAPT/KET/DOI in the DRN could dose-dependently shorten the time to awakening (P<0.01 or 0.01<P<0.05, Figure 5 F, H, I). Microinjection of WAY 100635 into the DRN resulted in prolonged recovery time (P<0.01, Figure 5 G). Among them, 3mg/mL, 6mg/mL 8-OH-DAPT and 2mg/ml DOI had a more significant reduction in awakening time (P<0.01, Figure 5F, H). The results were consistent with those of the pharmacological intervention section of the lateral ventricle. Combined with the results of the fourth part, it can be found that activation of 5-HT1A receptor can shorten the emergence time, while inhibition of 5-HT1A receptor can prolong the emergence time of sevoflurane anesthesia. However, activation of 5-HT2A/C receptor and inhibition of 5-HT2A receptor both led to shorter awakening times, which may be due to the different roles of different 5-HT receptors in awakening under general anesthesia. In addition, combined with the experimental results of the fourth part, we hypothesized that the stimulating effect of 5-HT1A receptor may play a leading role in regulating anesthesia recovery.

### 3.6 Activation of 5-HT1A receptor by injection of 8-OH-DAPT into the lateral ventricle (ICV) reduced the emergence time of sevoflurane anesthesia-(EEG)

In order to investigate the effects of sevoflurane anesthesia and 5-HT 1A receptor agonist 8-OH-DAPT on the brain electrical activity of mice, cortical EEG was recorded and analyzed under different conditions. We selected the same batch of wild-type C57BL/6J mice at 8 weeks after birth for lateral ventricle catheterization and cortical electrode implantation, 5-HT1A receptor agonist 8-OH-DAPT was administered through the lateral ventricle, and EEG was recorded. C57BL/6J mice were placed in the anesthesia box after one week of lateral ventricle catheterization, and the cortical EEG was recorded after the adaptation of the mice. After 5 minutes, the induction and maintenance of anesthesia began and the induction time was recorded. The 5-HT 1A receptor agonist 8-OH-DAPT 2000nL was administered 15 minutes after anesthesia began through a lateral ventricle catheter. The anesthesia was stopped at 30 minutes and the emergence time was recorded. Compared with the vehicle group, the induction time of sevoflurane anesthesia in the concentrations of 8-OH-DAPT (6mg/ml) group (FIG 6, P>0.05, n=8). Compared with the vehicle group, the emergence time of sevoflurane anesthesia in groups with in the concentrations of 8-OH-DAPT (6mg/ml) was significantly reduced (FIG 6, P<0.001, n=8). Cortical EEG of mice was recorded as anesthesia before sevo on, during sevo anesthesia and after sevo off (Figure 7A). We found that the cortical EEG voltage amplitude peak (power) decreased significantly after LORR in both groups, and the cortical EEG voltage was at a low level during the maintenance of anesthesia. When the mice were RORR, the cortical EEG voltage recovered, but the electrical activity was still lower than before anesthesia (Figure 7 B-F). For Delta wave and Theta wave, the average proportions of these three waves decreased significantly in the two groups after anesthesia compared with the state of being conscious, but the decrease of the average proportions in the 8-OH-DAPT group (0.01<P<0.05) was less than that in the vehicle group (P<0.01, Figure 6 G-H). For Alpha wave, the average proportion of Alpha wave in the vehicle group decreased significantly during anesthesia and in the awaking stage compared with that in the state of being conscious (P<0.01, FIG. 7-I). The average proportion of Alpha wave in the 8-OH-DAPT group decreased in the anesthesia stage but showed no statistical difference, when the mice were in the awaking stage, the proportion further decreased. Compared with the state of being conscious, there were significant differences (P<0.05, FIG. 7-I); For Beta waves, it in Vehicle group decreased significantly in anesthesia and awaking state compared with state of being conscious (P<0.01 or 0.01<P<0.05). However, in the 8-OH-DAPT group, there was no statistical difference in the ratio of Beta wave in anesthesia and awaking state compared with that in state of being conscious (P>0.05, FIG. 6-7). For Gamma wave, its proportion in vehicle group decreased after anesthesia but showed no statistical difference compared with the state of being conscious (P>0.05), but significantly decreased in the awaking state (P<0.01). For the 8-OH-DAPT group, the Gamma wave proportion decreased significantly under anesthesia and awaking state (P<0.05, FIG 7-K). Although there was no statistical difference in the proportion of each wave between the two groups in different states, it indicated that the injection of 8-OH-DAPT into the lateral ventricle had no significant effect on the depth of anesthesia. However, the amplitude of each wave in the 8-OH-DAPT group decreased less than that in the vehicle group during anesthesia and awaking state, indicating that the administration of 8-OH-DAPT in the lateral ventriculus can relieve the inhibition of brain electrical activity caused by anesthesia in mice, shorten the emergence time of anesthesia, and facilitate the recovery of mice after general anesthesia (FIG 6-7).

**Figure 6.**
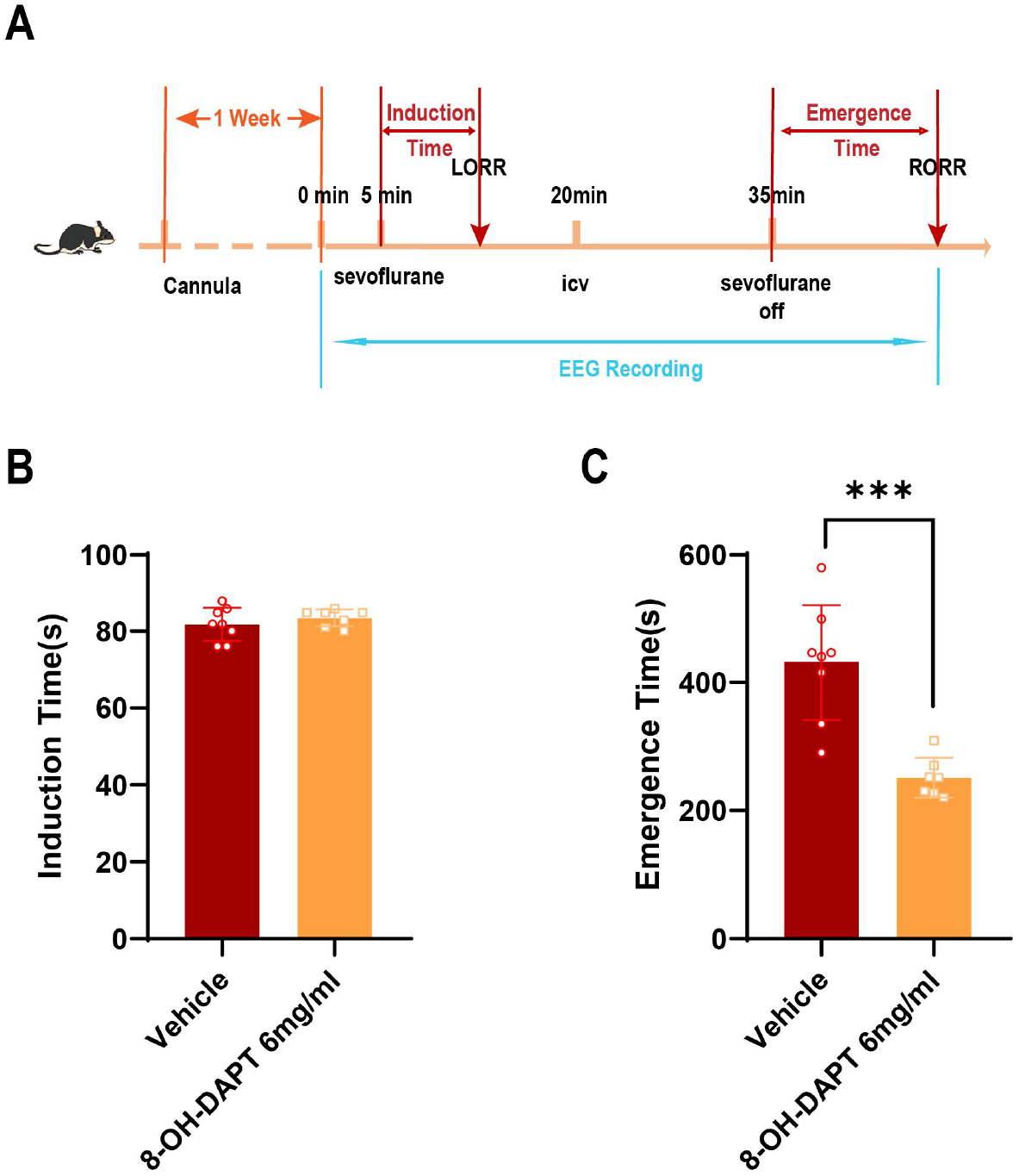
Effects of Vehicle/8-OH-DAPT administered in the lateral ventricle on EEG activity. A. Specific experimental protocols for EEG recording. (B) Compared with the vehicle group, the induction time of sevoflurane anesthesia in the concentrations of 8-OH-DAPT (6mg/ml) group (P>0.05, n=8). (C) Compared with the vehicle group, the emergence time of sevoflurane anesthesia in groups with in the concentrations of 8-OH-DAPT (6mg/ml) was significantly reduced.

**Figure 7.**
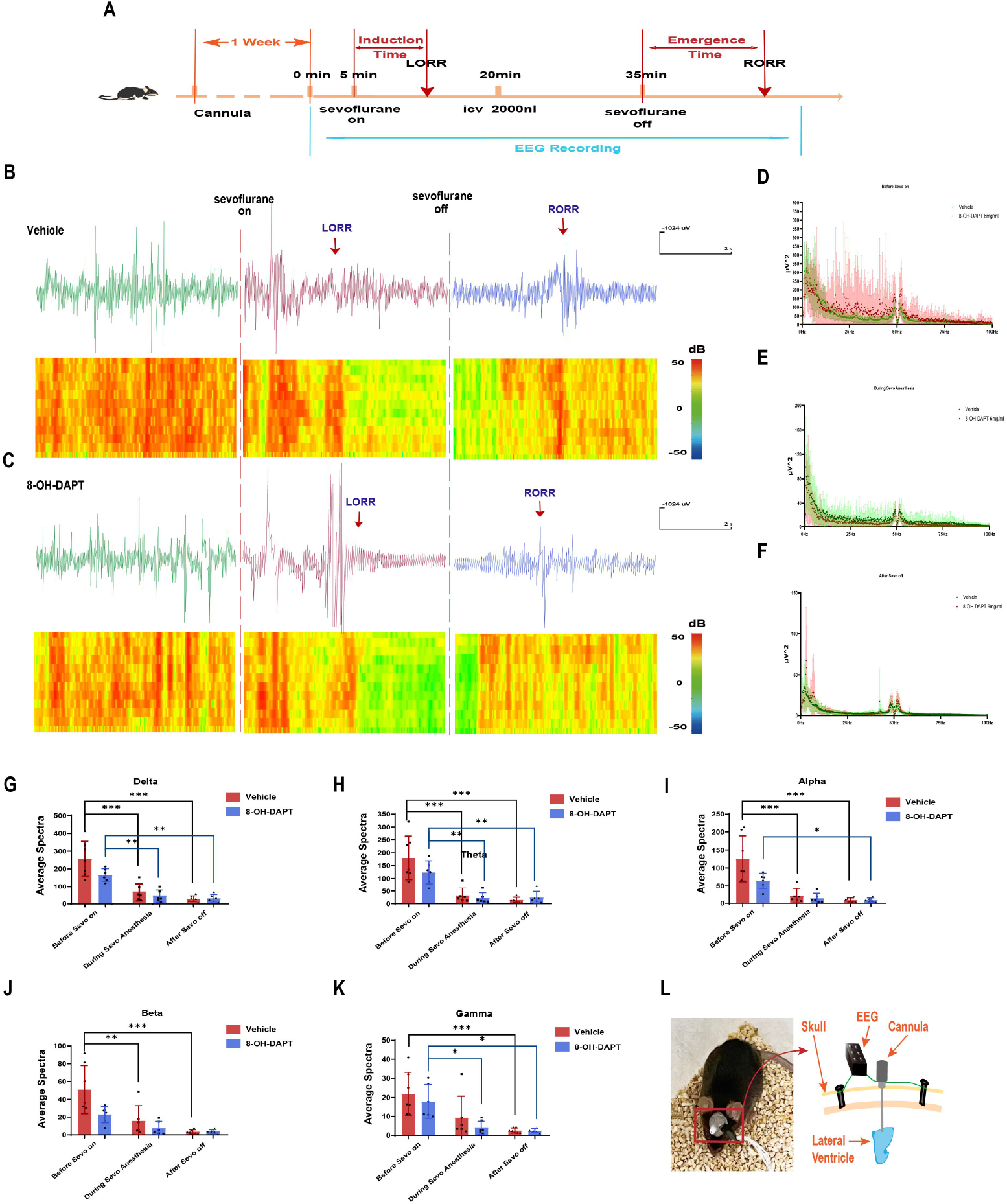
Effects of Vehicle/8-OH-DAPT administered in the lateral ventricle on EEG activity. (A) Specific experimental protocols for EEG recording. (B-C) EEG in vehicle and 8-OH-DAPT groups. (D-F) EEG power spectrum of state of being conscious, anesthetized and awakened rats (n=8). (G-K) The proportions of Delta, Theta, Alpha, Beta and Gamma waves in the two groups under different states. (L) Schematic diagram of cortical EEG recording and drug delivery device in the lateral ventricle.

## Discussion

In this study, 5-HT neurons in DRN are involved in the regulation of isoflurane anesthesia recovery through pharmacological and photogenetic experiments. Intraventricular infusion and intranuclear microinjection of serotonin and 5-HT receptor agonists or antagonists indicate that serotonin and 5-HT 1A and 2A/C receptor, especially 5-HT 1A receptor, are involved in the regulation of awakening delay mediated by DRN 5-HT neurons. Finally, we also observed the effect of anesthesia on EEG in mice by cortical EEG recording, indicating that activating 5-HT 1A receptor can alleviate the inhibition of EEG activity caused by anesthesia and shorten the recovery time of anesthesia.

The arousal of the brain depends on many regions that can receive different arousal signals at the same time, in which noradrenergic system[21], histaminergic system[22], 5-HT system, dopaminergic system[23], glutamatergic system and orexin system play an important role. Of course, anesthesia-arousal regulation is a complex network regulation process, and there are other unknown regulation pathways. The ascending reticular activing system plays a very essential role in maintaining the awakening state of the body. DRN is mainly rich in 5-HT neurons and the physiological role of 5-HT is extensive and the mechanism is complex. 5-HT binds to various receptors (comprising 7 gene families, with at least 14 distinct subtypes in mice, including both ionotropic and G protein-coupled receptors) that engage diverse signal transduction pathways[24–26], further increasing the complexity of the 5-HT system postsynaptically. As an important part of the ascending activation system, 5-HT plays an important role in maintaining arousal and alertness.

Several studies have reported that the concentration of 5-HT in prefrontal cortex and hypothalamus will be reduced during general anesthesia[27,12]. Several other studies have found that the activity of 5-HT neurons in DR is closely related to arousal behavior through multichannel recording. The level of 5-HT decreases significantly in slow wave sleep (SWS), while the activity of 5-HT neurons in DRN increases significantly when mice transition from sleep to arousal[28–30], which is consistent with the results of our study. However, recent researches suggest that the role of 5-HT in sleep and wake cannot be simply summarized as sleep-promoting or wakefulness-promoting. With the different time and degree of activation of central 5-HT system, the specific role of 5-HT on sleep or wake may vary and change from time to time[31–33]. These studies, combined with our findings, hint at the bidirectional regulatory role of 5-HT neurons in DRN and highlight the importance of further research to measure the activation of these neurons during wakefulness, sleep, or anesthesia.

Serotonin can bind to a variety of receptors with different characteristics, giving the serotonergic system the ability to regulate a variety of behavioral functions[15]. 5-HT 1A receptor is a metabolic receptor that can hyperpolarize membrane potential by activating G protein coupled receptor potassium channel[26]. Relevant studies have shown that the systematic application of 5-HT 1A receptor agonist flesinoxan or 8-OH-DPAT can increase arousal and reduce slow wave sleep (SWS) and rapid eye movement (REM) sleep[34]. Recently, the 5-HT1A receptor has been found to be expressed also by non-5-HT cells of the DRN[35,29]. Activation of postsynaptic 5-HT1A receptors expressed by GABAergic cells located in the DRN would be expected to indirectly facilitate the activity of 5-HT neurons. In this respect, Koyama et al. have shown that 5-HT reduced the frequency of miniature inhibitory postsynaptic currents arising from attached GABAergic presynaptic terminals recorded in dissociated rat basolateral amygdala nuclei[36]. For the 5-HT 2A/C receptor, The 5-HT2A and the 5-HT2C receptors have striking amino acid homology. They are primarily coupled to Gq protein and their actions are mediated by the activation of phospholipase C, with a resulting depolarization of the host cell. The serotonin 5-HT2A and 5-HT2C receptor-containing neurons are predominantly GABAergic interneurons and projection neurons[37–40]. Thus, activation of 5-HT2A and 5-HT2C receptors expressed by GABAergic cells located in the DRN would result in the decrease 5-HT neuronal firing rate. Therefore, the presence of 5-HT receptors within the boundaries of the DRN, expressed by GABAergic interneurons, tends to indicate the existence of a supplemental mechanism in the control of 5-HT neurons functional activity.

In this study, 5-HT 1A receptor agonists or antagonists were injected into the lateral ventricle and DRN. It was found that activating 5-HT 1A receptor can significantly shorten the recovery time of sevoflurane anesthesia. Combined with this study, the conclusions of our research supports the important role of 5-HT 1A receptor in the recovery from general anesthesia. For the 5-HT 2A/C receptor, some studies have shown that the activation of serotonergic neurons may partially promote the awakening of general anesthesia through the 5-HT 2C receptor[41]. Through intracerebroventricular injection of 5-HT 2A/C receptor agonist DOI and 5-HT 2C receptor antagonist RS-102221, it was found that activating 5-HT 2A/C receptor can shorten the awakening time and inhibiting 5-HT 2C receptor can prolong the awakening time. Therefore, they speculated that it was the antagonist of 5-HT 2C receptor rather than the antagonist of 5-HT 2A receptor that had the opposite effect to DOI. In this study, we found that injection of 5-HT 2A/C receptor agonist DOI into lateral ventricle and DRN can significantly shorten the recovery time, while injection of 5-HT 2A receptor antagonist KET can also shorten the recovery time. Combined with the above conclusions, this study supplemented the role of 5-HT 2A receptor in anesthesia awakening: 5-HT 2A receptor has an inhibitory effect in general anesthesia awakening. This further shows that 5-HT system can play different roles in general anesthesia awakening through different receptors. Combined with our experimental results of photogenetic activation of 5-HT neurons and exogenous supplement of 5-HT, we speculate that 5-HT 1A receptor plays a major and key role in promoting the recovery from general anesthesia.

In order to clarify the mechanism of 5-HT neurons in general anesthesia, we observed the activity of 5-HT DRN neurons by testing the c-fos expressed in the TPH2-neurons under general anesthesia and the activity of 5-HT DRN neurons can be strongly suppressed by sevoflurane anesthesia. Subsequently,we used EEG recording to record the cortical EEG of mice in different states and, to some extent, EEG can reflect the depth of anesthesia. Our experiment found that sevoflurane anesthesia can significantly inhibit the cortical EEG activity of mice. Although compared with vehicle control group, there was no significant difference in the cortical EEG activity of 8-OH-DAPT injected into the lateral ventricle under anesthesia and awakening, the application of 8-OH-DAPT can alleviate the inhibition of EEG activity in mice caused by anesthesia, promote the recovery of mice after general anesthesia and shorten the recovery time of anesthesia. This effect of promoting recovery may not be achieved by changing the depth of anesthesia, but by other ways. Combined with the previous research findings of our research group: in DBA/1 mice, activating 5-HT neurons in DR can reduce the incidence of sudden epileptic death by reducing epilepsy induced respiratory arrest (S-IRA)[42,43]. Other study indicates that developmental loss of Pet1 5-HT neurons, through Pet1-Cre-driven deletion of Lmx1b, leads to blunting of the chemorespiratory reflex in adult mice[44]. Compared with controls, these mutant Lmx1bf/f/p mice showed a diminished ventilatory response to hypercapnia[45]. These suggest that 5-HT neurons in DR play an important role in regulating respiration. Therefore, we further speculate that 5-HT in DR can promote the recovery of general anesthesia by regulating respiration rather than changing the depth of anesthesia. Nevertheless, its specific mechanism needs to be further studied.

### The limitations to this study

Although we have experimentally verified the role of 5-HT 1A, 2A/C and 2A receptors in awakening under general anesthesia, there are many other 5-HT receptors except these receptors, and the role of these receptors in awakening under general anesthesia needs to be further studied. Secondly, we verified the involvement of 5-HT in the recovery from general anesthesia through drug administration in the lateral ventricle and the DR nucleus, but it could not be ruled out that activation of 5-HT in DRN might also release glutamate and other neurotransmitters. Thirdly, whether the activation of 5-HT neurons in DR is generated by regulating respiration also needs to be further verified.

## CRediT authorship contribution statement

HaiXiang Ma, LeYuan Gu, YuLing Wang, Qing Xu, Qian Yu and XiTing Lian perfromed the experiments. Lu Liu, JiaXuan Gu 1, WeiHui Shao1, Yue Shen contributed to the formal analysis. YuLing Wang and Qing Xu contributed to writing the paper. HongHai Zhang designed the experiment and wrote the paper.

## Reference

[1] Brown E N, Purdon P L, Van Dort C J. General anesthesia and altered states of arousal: a systems neuroscience analysis[J/OL]. Annual Review of Neuroscience, 2011, 34: 601–628. DOI:10.1146/annurev-neuro-060909-153200.

[2] Franks N P. General anaesthesia: from molecular targets to neuronal pathways of sleep and arousal[J/OL]. Nature Reviews. Neuroscience, 2008, 9(5): 370–386. DOI:10.1038/nrn2372.

[3] Makaryus R, Lee H, Yu M, et al. The metabolomic profile during isoflurane anesthesia differs from propofol anesthesia in the live rodent brain[J/OL]. Journal of Cerebral Blood Flow and Metabolism: Official Journal of the International Society of Cerebral Blood Flow and Metabolism, 2011, 31(6): 1432–1442. DOI:10.1038/jcbfm.2011.1.

[4] Xie H, Chung D Y, Kura S, et al. Differential effects of anesthetics on resting state functional connectivity in the mouse[J/OL]. Journal of Cerebral Blood Flow & Metabolism, 2020, 40(4): 875–884. DOI:10.1177/0271678X19847123.

[5] Harris M, Chung F. Complications of general anesthesia[J/OL]. Clinics in Plastic Surgery, 2013, 40(4): 503–513. DOI:10.1016/j.cps.2013.07.001.

[6] Biccard B M, Rodseth R N. The pathophysiology of peri-operative myocardial infarction: Pathophysiology of peri-operative myocardial infarction[J/OL]. Anaesthesia, 2010, 65(7): 733–741. DOI:10.1111/j.1365-2044.2010.06338.x.

[7] Evered L, Scott D A, Silbert B, et al. Postoperative cognitive dysfunction is independent of type of surgery and anesthetic[J/OL]. Anesthesia and Analgesia, 2011, 112(5): 1179–1185. DOI:10.1213/ANE.0b013e318215217e.

[8] Smetana G W. Preoperative pulmonary evaluation[J/OL]. The New England Journal of Medicine, 1999, 340(12): 937–944. DOI:10.1056/NEJM199903253401207.

[9] Ferreyra G, Long Y, Ranieri V M. Respiratory complications after major surgery[J/OL]. Current Opinion in Critical Care, 2009, 15(4): 342–348. DOI:10.1097/MCC.0b013e32832e0669.

[10] Leslie K, Chan M T V, Myles P S, et al. Posttraumatic stress disorder in aware patients from the B-aware trial[J/OL]. Anesthesia and Analgesia, 2010, 110(3): 823–828. DOI:10.1213/ANE.0b013e3181b8b6ca.

[11] Harper N J N, Dixon T, DuguÉ P, et al. Suspected anaphylactic reactions associated with anaesthesia[J/OL]. Anaesthesia, 2009, 64(2): 199–211. DOI:10.1111/j.1365-2044.2008.05733.x.

[12] Mukaida K, Shichino T, Koyanagi S, et al. Activity of the serotonergic system during isoflurane anesthesia[J/OL]. Anesthesia and Analgesia, 2007, 104(4): 836–839. DOI:10.1213/01.ane.0000255200.42574.22.

[13] Roizen M F, White P F, Eger E I, et al. Effects of ablation of serotonin or norepinephrine brain-stem areas on halothane and cyclopropane MACs in rats[J/OL]. Anesthesiology, 1978, 49(4): 252–255. DOI:10.1097/00000542-197810000-00005.

[14] Monti J M. Serotonin control of sleep-wake behavior[J/OL]. Sleep Medicine Reviews, 2011, 15(4): 269–281. DOI:10.1016/j.smrv.2010.11.003.

[15] Okaty B W, Commons K G, Dymecki S M. Embracing diversity in the 5-HT neuronal system[J/OL]. Nature Reviews Neuroscience, 2019, 20(7): 397–424. DOI:10.1038/s41583-019-0151-3.

[16] Monti J M. The structure of the dorsal raphe nucleus and its relevance to the regulation of sleep and wakefulness[J/OL]. Sleep Medicine Reviews, 2010, 14(5): 307–317. DOI:10.1016/j.smrv.2009.11.004.

[17] Ren J, Friedmann D, Xiong J, et al. Anatomically Defined and Functionally Distinct Dorsal Raphe Serotonin Sub-systems[J/OL]. Cell, 2018, 175(2): 472-487.e20. DOI:10.1016/j.cell.2018.07.043.

[18] Mohr A A, Garcia-Serrano A M, Vieira J P, et al. A glucose-stimulated BOLD fMRI study of hypothalamic dysfunction in mice fed a high-fat and high-sucrose diet[J/OL]. Journal of Cerebral Blood Flow and Metabolism: Official Journal of the International Society of Cerebral Blood Flow and Metabolism, 2021, 41(7): 1734–1743. DOI:10.1177/0271678X20942397.

[19] Hopkins S C, Dedic N, Koblan K S. Effect of TAAR1/5-HT1A agonist SEP-363856 on REM sleep in humans[J/OL]. Translational Psychiatry, 2021, 11(1): 228. DOI:10.1038/s41398-021-01331-9.

[20] Yelleswarapu N K, Tay J K, Fryer W M, et al. Elucidating the role of 5-HT1A and 5-HT7 receptors on 8-OH-DPAT-induced behavioral recovery after experimental traumatic brain injury[J/OL]. Neuroscience Letters, 2012, 515(2): 153–156. DOI:10.1016/j.neulet.2012.03.033.

[21] Moore J T, Chen J, Han B, et al. Direct Activation of Sleep-Promoting VLPO Neurons by Volatile Anesthetics Contributes to Anesthetic Hypnosis[J/OL]. Current Biology, 2012, 22(21): 2008–2016. DOI:10.1016/j.cub.2012.08.042.

[22] Yamanaka A, Tsujino N, Funahashi H, et al. Orexins activate histaminergic neurons via the orexin 2 receptor[J/OL]. Biochemical and Biophysical Research Communications, 2002, 290(4): 1237–1245. DOI:10.1006/bbrc.2001.6318.

[23] Eriksson K S, Sergeeva O, Brown R E, et al. Orexin/hypocretin excites the histaminergic neurons of the tuberomammillary nucleus[J]. The Journal of Neuroscience: The Official Journal of the Society for Neuroscience, 2001, 21(23): 9273–9279.

[24] Hannon J, Hoyer D. Molecular biology of 5-HT receptors[J/OL]. Behavioural Brain Research, 2008, 195(1): 198–213. DOI:10.1016/j.bbr.2008.03.020.

[25] Hoyer D. 5-HT Receptor Nomenclature: Naming Names, Does It Matter? A Tribute to Maurice Rapport[J/OL]. ACS chemical neuroscience, 2017, 8(5): 908–919. DOI:10.1021/acschemneuro.7b00011.

[26] Filip M, Bader M. Overview on 5-HT receptors and their role in physiology and pathology of the central nervous system[J/OL]. Pharmacological reports: PR, 2009, 61(5): 761–777. DOI:10.1016/s1734-1140(09)70132-x.

[27] Fiske E, Portas C M, GrØNli J, et al. Increased extracellular 5-HT but no change in sleep after perfusion of a 5-HT1A antagonist into the dorsal raphe nucleus of rats[J/OL]. Acta Physiologica (Oxford, England), 2008, 193(1): 89–97. DOI:10.1111/j.1748-1716.2007.01792.x.

[28] Oikonomou G, Altermatt M, Zhang R W, et al. The Serotonergic Raphe Promote Sleep in Zebrafish and Mice[J/OL]. Neuron, 2019, 103(4): 686-701.e8. DOI:10.1016/j.neuron.2019.05.038.

[29] Kirby L G, Pernar L, Valentino R J, et al. Distinguishing characteristics of serotonin and non-serotonin-containing cells in the dorsal raphe nucleus: electrophysiological and immunohistochemical studies[J/OL]. Neuroscience, 2003, 116(3): 669–683. DOI:10.1016/S0306-4522(02)00584-5.

[30] Trulson M E, Jacobs B L. Raphe unit activity in freely moving cats: correlation with level of behavioral arousal[J/OL]. Brain Research, 1979, 163(1): 135–150. DOI:10.1016/0006-8993(79)90157-4.

[31] Morrow J D, Vikraman S, Imeri L, et al. Effects of serotonergic activation by 5-hydroxytryptophan on sleep and body temperature of C57BL/6J and interleukin-6-deficient mice are dose and time related[J/OL]. Sleep, 2008, 31(1): 21–33. DOI:10.1093/sleep/31.1.21.

[32] Linthorst A C E, Flachskamm C, Hopkins S J, et al. Long-Term Intracerebroventricular Infusion of Corticotropin-Releasing Hormone Alters Neuroendocrine, Neurochemical, Autonomic, Behavioral, and Cytokine Responses to a Systemic Inflammatory Challenge[J/OL]. The Journal of Neuroscience, 1997, 17(11): 4448–4460. DOI:10.1523/JNEUROSCI.17-11-04448.1997.

[33] Imeri L, Mancia M, Bianchi S, et al. 5-Hydroxytryptophan, but not L-tryptophan, alters sleep and brain temperature in rats[J/OL]. Neuroscience, 2000, 95(2): 445–452. DOI:10.1016/s0306-4522(99)00435-2.

[34] MONTI null, JANTOS null. Dose-dependent effects of the 5-HT1A receptor agonist 8-OH-DPAT on sleep and wakefulness in the rat[J/OL]. Journal of Sleep Research, 1992, 1(3): 169–175. DOI:10.1111/j.1365-2869.1992.tb00033.x.

[35] Day H E W, Greenwood B N, Hammack S E, et al. Differential expression of 5HT-1A, alpha 1b adrenergic, CRF-R1, and CRF-R2 receptor mRNA in serotonergic, gamma-aminobutyric acidergic, and catecholaminergic cells of the rat dorsal raphe nucleus[J/OL]. The Journal of Comparative Neurology, 2004, 474(3): 364–378. DOI:10.1002/cne.20138.

[36] Koyama S, Kubo C, Rhee J, et al. Presynaptic serotonergic inhibition of GABAergic synaptic transmission in mechanically dissociated rat basolateral amygdala neurons[J/OL]. The Journal of Physiology, 1999, 518(2): 525–538. DOI:10.1111/j.1469-7793.1999.0525p.x.

[37] Clemett D A, Punhani T, Duxon M S, et al. Immunohistochemical localisation of the 5-HT2C receptor protein in the rat CNS[J/OL]. Neuropharmacology, 2000, 39(1): 123–132. DOI:10.1016/s0028-3908(99)00086-6.

[38] LÓPez-GimÉNez J F, VilarÓ M T, Palacios J M, et al. Mapping of 5-HT2A receptors and their mRNA in monkey brain: [3H]MDL100,907 autoradiography and in situ hybridization studies[J/OL]. The Journal of Comparative Neurology, 2001, 429(4): 571–589. DOI:10.1002/1096-9861(20010122)429:4<571::aid-cne5>3.0.co;2-x.

[39] Mirkes S J, Bethea C L. Oestrogen, progesterone and serotonin converge on GABAergic neurones in the monkey hypothalamus[J/OL]. Journal of Neuroendocrinology, 2001, 13(2): 182–192. DOI:10.1046/j.1365-2826.2001.00612.x.

[40] Serrats J, Mengod G, CortÉs R. Expression of serotonin 5-HT2C receptors in GABAergic cells of the anterior raphe nuclei[J/OL]. Journal of Chemical Neuroanatomy, 2005, 29(2): 83–91. DOI:10.1016/j.jchemneu.2004.03.010.

[41] Li A, Li R, Ouyang P, et al. Dorsal raphe serotonergic neurons promote arousal from isoflurane anesthesia[J/OL]. CNS Neuroscience & Therapeutics, 2021, 27(8): 941–950. DOI:10.1111/cns.13656.

[42] Zhang H, Zhao H, Yang X, et al. 5-Hydroxytryptophan, a precursor for serotonin synthesis, reduces seizure-induced respiratory arrest[J/OL]. Epilepsia, 2016, 57(8): 1228–1235. DOI:10.1111/epi.13430.

[43] Zhang H, Zhao H, Zeng C, et al. Optogenetic activation of 5-HT neurons in the dorsal raphe suppresses seizure-induced respiratory arrest and produces anticonvulsant effect in the DBA/1 mouse SUDEP model[J/OL]. Neurobiology of Disease, 2018, 110: 47–58. DOI:10.1016/j.nbd.2017.11.003.

[44] Richardson-Jones J W, Craige C P, Nguyen T H, et al. Serotonin-1A autoreceptors are necessary and sufficient for the normal formation of circuits underlying innate anxiety[J/OL]. The Journal of Neuroscience: The Official Journal of the Society for Neuroscience, 2011, 31(16): 6008–6018. DOI:10.1523/JNEUROSCI.5836-10.2011.

[45] Hodges M R, Tattersall G J, Harris M B, et al. Defects in breathing and thermoregulation in mice with near-complete absence of central serotonin neurons[J/OL]. The Journal of Neuroscience: The Official Journal of the Society for Neuroscience, 2008, 28(10): 2495–2505. DOI:10.1523/JNEUROSCI.4729-07.2008.

